# Potential Rhodopsin and Bacteriochlorophyll-Based Dual Phototrophy in a High Arctic Glacier

**DOI:** 10.1101/2020.09.28.316679

**Authors:** Yonghui Zeng, Xihan Chen, Anne Mette Madsen, Athanasios Zervas, Tue Kjærgaard Nielsen, Adrian-Stefan Andrei, Lars Chresten Lund-Hansen, Yongqin Liu, Lars Hestbjerg Hansen

**Author notes:** Correspondence: Yonghui Zeng, Department of Environmental Science, Aarhus University, Frederiksborgvej 399, Roskilde 4000, Denmark. or.

## Abstract

Conserving additional energy from sunlight through bacteriochlorophyll (BChl)‐based reaction center or proton‐pumping rhodopsin is a highly successful life strategy in environmental bacteria. Rhodopsin and BChl based systems display contrasting characteristics in the size of coding operon, cost of biosynthesis, ease of expression control, and efficiency of energy production. This raises an intriguing question of whether a single bacterium has evolved the ability to perform these two types of phototrophy complementarily according to energy needs and environmental conditions. Here we report four *Tardiphaga* sp. strains (Alphaproteobacteria) of monophyletic origin isolated from a high Arctic glacier in northeast Greenland (81.566° N, 16.363° W) that are at different evolutionary stages concerning phototrophy. Their >99.8% identical genomes contain footprints of horizontal operon transfers (HOT) of the complete gene clusters encoding BChl and xanthorhodopsin (XR)‐based dual phototrophy. Two strains only possess a complete xanthorhodopsin (XR) operon, while the other two strains have both a photosynthesis gene cluster (PGC) and an XR operon in their genomes. All XR operons are heavily surrounded by mobile genetic elements and located close to a tRNA gene, strongly signaling that a HOT event of XR operon has occurred recently. Mining public genome databases and our High Arctic glacial and soil metagenomes revealed that phylogenetically diverse bacteria have the metabolic potential of performing BChl and rhodopsin‐based dual phototrophy. Our data provide new insights on how bacteria cope with the harsh and energy‐deficient environments in surface glaciers, possibly by maximizing the capability of exploiting solar energy.

**Importance:** Over billions of years of evolution, bacteria capable of light‐driven energy production have occupied every corner of surface Earth where solar irradiation can reach. Only two general biological systems have evolved in bacteria to be capable of net energy conservation via light‐harvesting: one is based on the pigment of (bacterio‐)chlorophyll and the other based on light‐sensing retinal molecules. There is emerging genomic evidence that these two rather different systems can co‐exist in a single bacterium to take advantage of their contrasting characteristics in the number of genes involved, biosynthesis cost, ease of expression control and efficiency of energy production, and thus enhance the capability of exploiting solar energy. Our data provide the first clear‐cut evidence that such dual phototrophy potentially exist in glacial bacteria. Further public genome mining suggests this understudied dual phototrophic mechanism is possibly more common than our data alone suggested.

**Sequence data availability:** Genomes, metagenomes and raw reads were deposited into GenBank under Bioprojects PRJNA548505 and PRJNA552582.

## Main Text

Over billions of years of evolution, phototrophic bacteria capable of light‐driven energy generation have occupied every corner of surface Earth where solar irradiation can reach. Only two general biological systems are known in bacteria to be capable of net energy conservation from light harvesting: one is based on bacteriochlorophyll (BChl; chlorophyll in the case of Cyanobacteria) and the other based on retinal molecules (1). BChl‐based system relies on a complex system consisting of dozens of proteins and pigments to form reaction center and antenna complex. In contrast, rhodopsin‐based system only requires a few genes to operate, including a key pair of genes, the rhodopsin gene and the carotenoid oxygenase gene *blh/brp* for retinal biosynthesis (2), albeit in a much lower efficiency in energy production than BChl‐ based system (3).

Rhodopsin and BChl based systems display contrasting characteristics in the size of coding operons, cost of biosynthesis, ease of expression control and efficiency of energy production. This raises an intriguing question of whether a single bacterium can employ both types of phototrophy to take advantage of their complementary properties in order to increase the flexibility in energy production. Given the high abundance and frequent co‐existence of BChl‐based phototrophs and rhodopsin‐based phototrophs in the same environment, for instance, oceans (4), and that phototrophy related genes frequently occurred in extrachromosomal genetic elements, like PGC on plasmids (5,6) or chromid (7), and proteorhodopsin in viral genomes (8), BChl and rhodopsin‐based dual phototrophic bacteria very likely have evolved in nature, awaiting discovery.

Indeed, there is emerging genomic evidence for such dual phototrophy. Recently, three *Roseiflexus* genomes (Chloroflexi phylum) from spring microbial mats were found to contain both *pufM* (encoding the M subunit of RC) and xanthorhodopsin (XR)‐ like gene (9), including two metagenome‐assembled genomes (MAGs) of *Roseiflexus* spp. OTU‐1 and OTU‐6 (10) and one from the isolate *Roseiflexus* sp. RS‐1 (11), albeit it is unclear if their rhodopsins function as bona fide proton pump, owing to the missing of the key carotenoid oxygenase gene *blp/brh* that is often located in the neighborhood of rhodopsin gene (10).

### Discovery of glacial bacteria with potential dual phototrophy

Aiming to provide further pure culture and direct evolutionary evidence for such potential dual phototrophy, we conducted both cultivation and metagenomics survey in the “Lille Firn” glacier (LF) and nearby exposed soil (ES) in northeast Greenland (81.566° N, 16.363° W; Fig. S1). A collection of isolates of aerobic phototrophic bacteria was created from the LF surface glacial ice sample. Four pinkish colonies (strains vice154, vice278, vice304, and vice352) were further examined due to their high similarities in matrix‐assisted laser desorption/ionization‐time of flight mass spectrometry profiling and the observation that vice154 and vice278 displayed weak BChl fluorescence signals using an infrared imaging technique (12). The 16S rRNA genes of the four strains are 100% identical and share 96.5% identity to *T. robiniae* LMG 26467^T^ of the genus *Tardiphaga* of Alphaproteobacteria (13), representing a novel species in the cryosphere cluster of *Tardiphaga* (Fig. 1A). In the glacial bacterial community (LF), members of *Tardiphaga* accounted for 0.017% (17/101,183) (Fig. S2). No *Tardiphaga*‐affiliated read was found in ES (n=10,060). *Tardiphaga* represents one of the least abundant groups and only occurs in LF.

**Figure 1.**
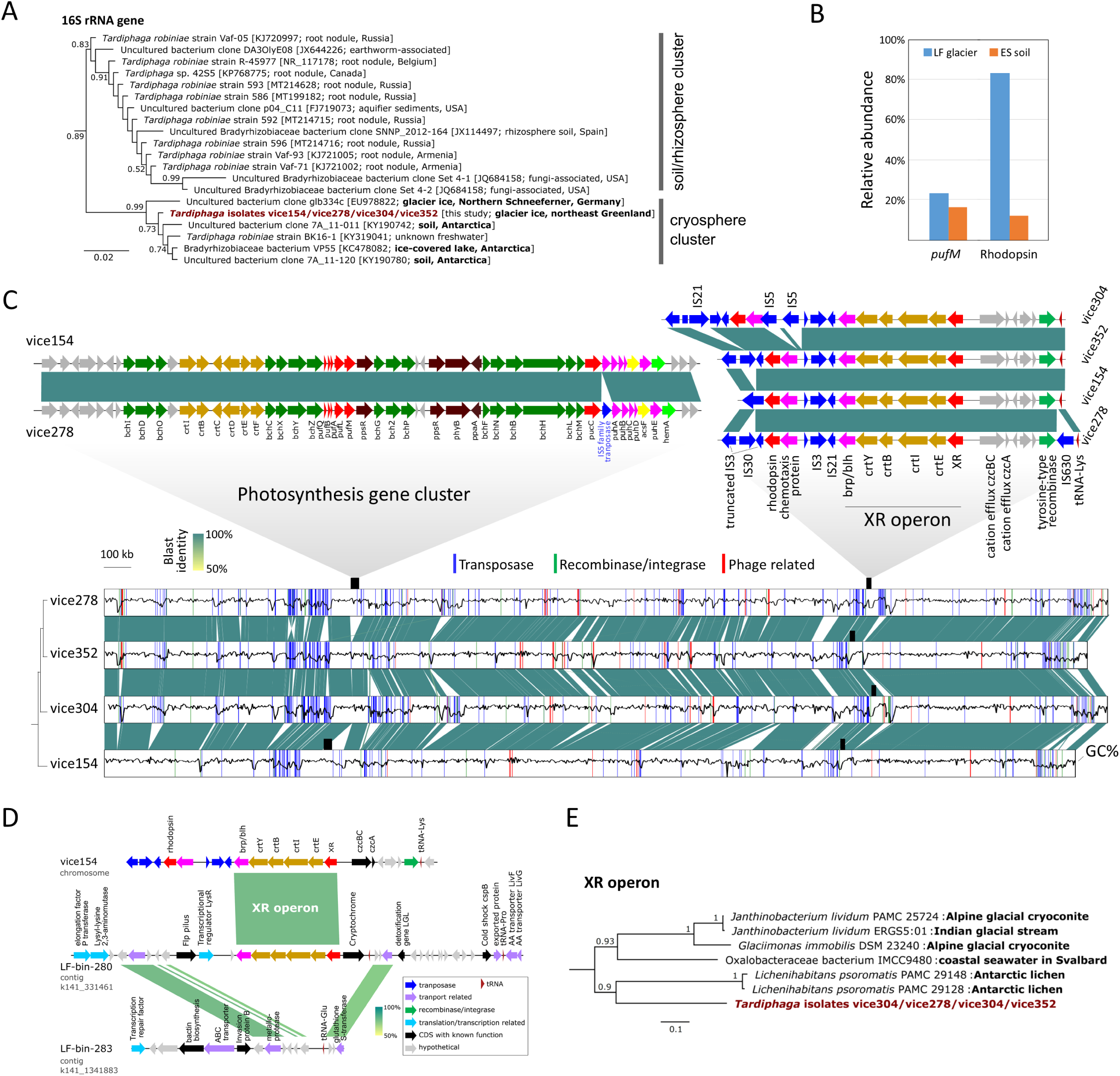
(**A**), maximum‐likelihood phylogeny of 16S rRNA genes. (**B**), abundance of *pufM* and rhodopsin genes in the LF and ES metagenomes, calculated as the number of mapped reads normalized to per kb length divided by the normalized number of reads mapped to the housekeeping gene *recA*. (**C**), genome synteny and sequence similarities of *Tardiphaga* sp. strains vice154, vice278, vice304, and vice352. GenBank accession number and source environment of each 16S rRNA gene sequence are shown in brackets on the tree. The architecture of photosynthesis gene cluster (PGC) and xanthorhodopsin (XR) operon are highlighted. Gaps in the alignment show non‐ conserved regions caused by mobilome related activities, including transposase, recombinase, integrase or phage‐related genes, the location of which are highlighted in the genome. All genomes are complete and start at the replication origin loci. Black wave lines inside the genomes (shown as box regions) represent the variations in GC content calculated in a window of 1 kb. The relationship between strains were estimated as a distance‐based genome tree in the bottom left using MASH (https://github.com/marbl/Mash). Color bar, BLAST identities. (**D**), genome synteny between contigs in the XR‐bearing glacial *Tardiphaga* MAG LF‐bin‐280 and non‐ phototrophic glacial *Tardiphaga* MAG LF‐bin‐283 in reference to the genome of strain vice154. (**E**), maximum‐likelihood phylogeny of the whole XR operon of *Tardiphaga* isolates and their top tBLASTn hits in NCBI’s genome database, for the full version of the tree see Fig. S5.

Despite their monophyletic origin as reflected by >99.8% genome pairwise average nucleotide identities and the highly conserved genome synteny (Fig. 1C), these four strains differed in both genome size and GC content (Table S1). These differences were primarily caused by insertions and deletions (Fig. 1C), including a 45.7‐kb PGC that is present in vice154 and vice278 but absent in vice304 and vice352. These two PGCs only differ in an insertion of an IS*5* family transposase gene between *pucC* and *puhA* in vice278 (Fig. 1C). No mobilome‐related genes were found in the proximity of PGC in both vice154 and vice278 (Fig. S3), suggesting that PGC is an ancient trait in the ancestor that later lost in vice304 and vice352.

All four genomes contain an XR operon encoding XR‐based phototrophy with the same gene arrangement (*XR‐crtEIBY‐brp*) and identical sequences (only one base difference out of 6,257 sites occurring in vice304) (Fig. 1C). The XR sequence contains most of the conserved sites including key residues as proton acceptor and donor (Fig. S4), suggesting that it very likely encodes a functional proton pump. Interestingly, there are an additional unclassified rhodopsin gene and a putative methyl‐accepting chemotaxis protein gene located upstream of the XR operon and flanked by insertion (IS) elements at both sides.

The XR operon is located near the tRNA‐Lys gene, a signal of recombination hotspot (14). The existence of multiple IS elements and a tRNA gene surrounding the XR operon strongly indicates that a HOT event of XR operon has occurred in these four strains. This was further supported by the reconstruction of two *Tardiphaga* MAGs (LF‐bin‐280 with the XR operon and LF‐bin‐283 without the XR operon; Table 1), where HOT of a complete XR operon was recorded at a highly homologous region next to a tRNA gene (Fig. 1D). Since the first discovery of proteorhodopsins in the oceans (15), there is a growing body of phylogeny‐based evidence for horizontal transfer of rhodopsin genes occurring between prokaryotes (16,17,18). Our data provide further clear‐cut evidence for HOT of a rhodopsin operon occurring in a natural community.

**Table 1.**
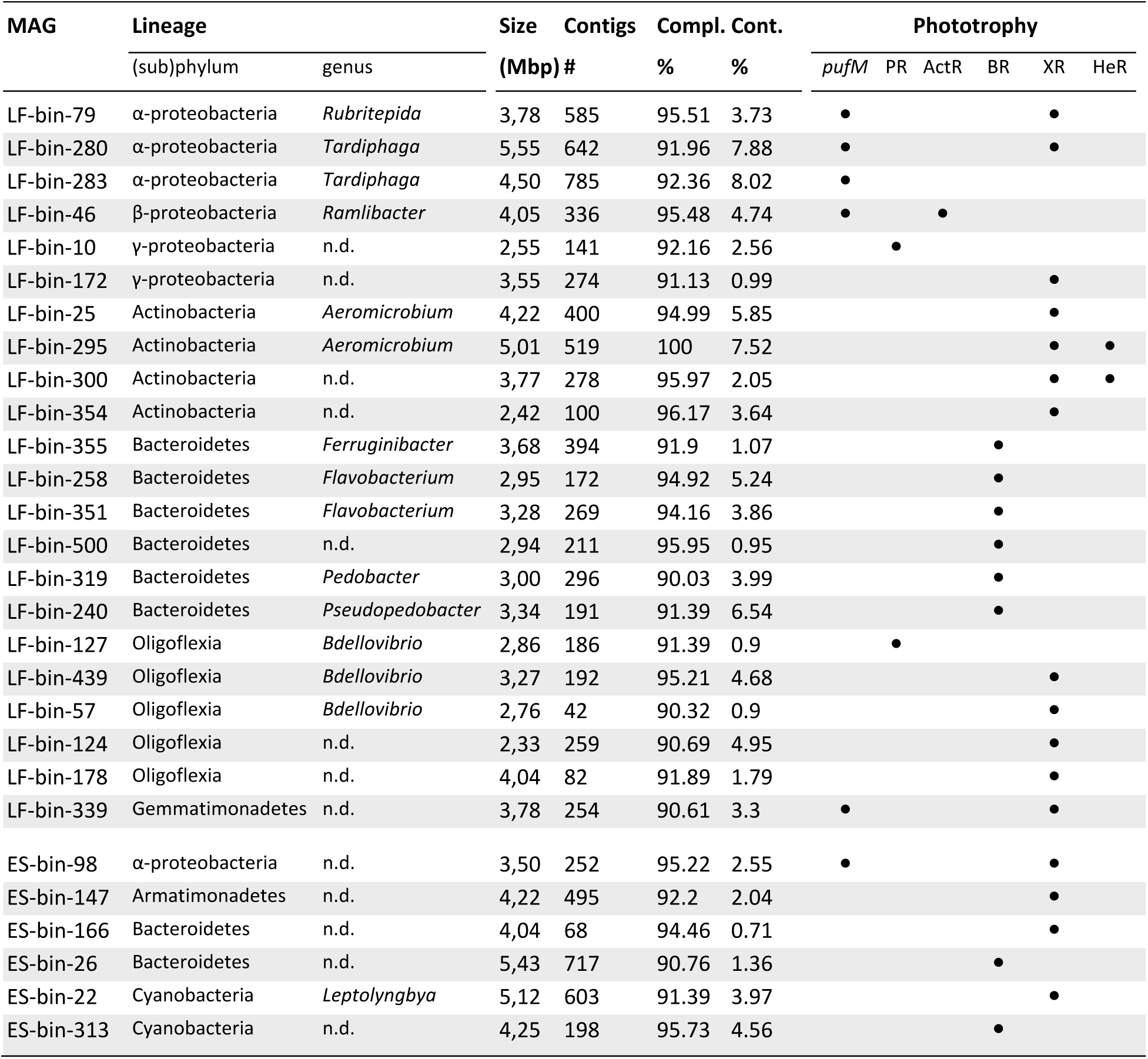
Summary of metagenome‐assembled genomes (MAGs) reconstructed from the metagenomes of the “Little Firn” glacier (LF) and nearby exposed soil (ES) that contain genes related to bacteriochlorophyll or rhodopsin‐based phototrophy. Only MAGs that are >90% complete with <10% contamination are shown. See the Table S2 for the full list of MAGs (>50% complete and <10% contamination). PR, proteorhodopsin; XR, xanthorhodopsin; ActR, actinorhodopsin; BR, bacteriorhodopsin; HeR, heliorhodopsin. n.d., not determined. Classification of rhodopsins were based on phylogenetic analysis with a comprehensive collection of rhodopsin genes as references (see Supplementary Methods).

The HOT of XR operon was likely driven by transposon activities as indicated by the presence of an integrase gene of the tyrosine‐type recombinase family and direct repeats or inverted repeats that can serve as attachment sites for recombinase on the XR transposon and IS*630* composite transposon predicted bioinformatically (Fig. 2A). The putative XR transposon is conserved and thus may occur in all *Tardiphaga* strains. Strain vice278 possesses an additional IS*630* family transposase between XR operon and tRNA, which has an identical copy 45.7 kb apart upstream of the XR operon (Fig. 2A). The two identical IS*630* genes can serve as IR for forming the putative IS*630* composite transposon through the DR of IRL IS*630* or IRR IS*630*. Given the almost 100% identical sequence of the whole XR operon in four *Tardiphaga* genomes, the acquisition of an XR operon in the ancestor certainly occurred before PGC divergence.

**Figure 2.**
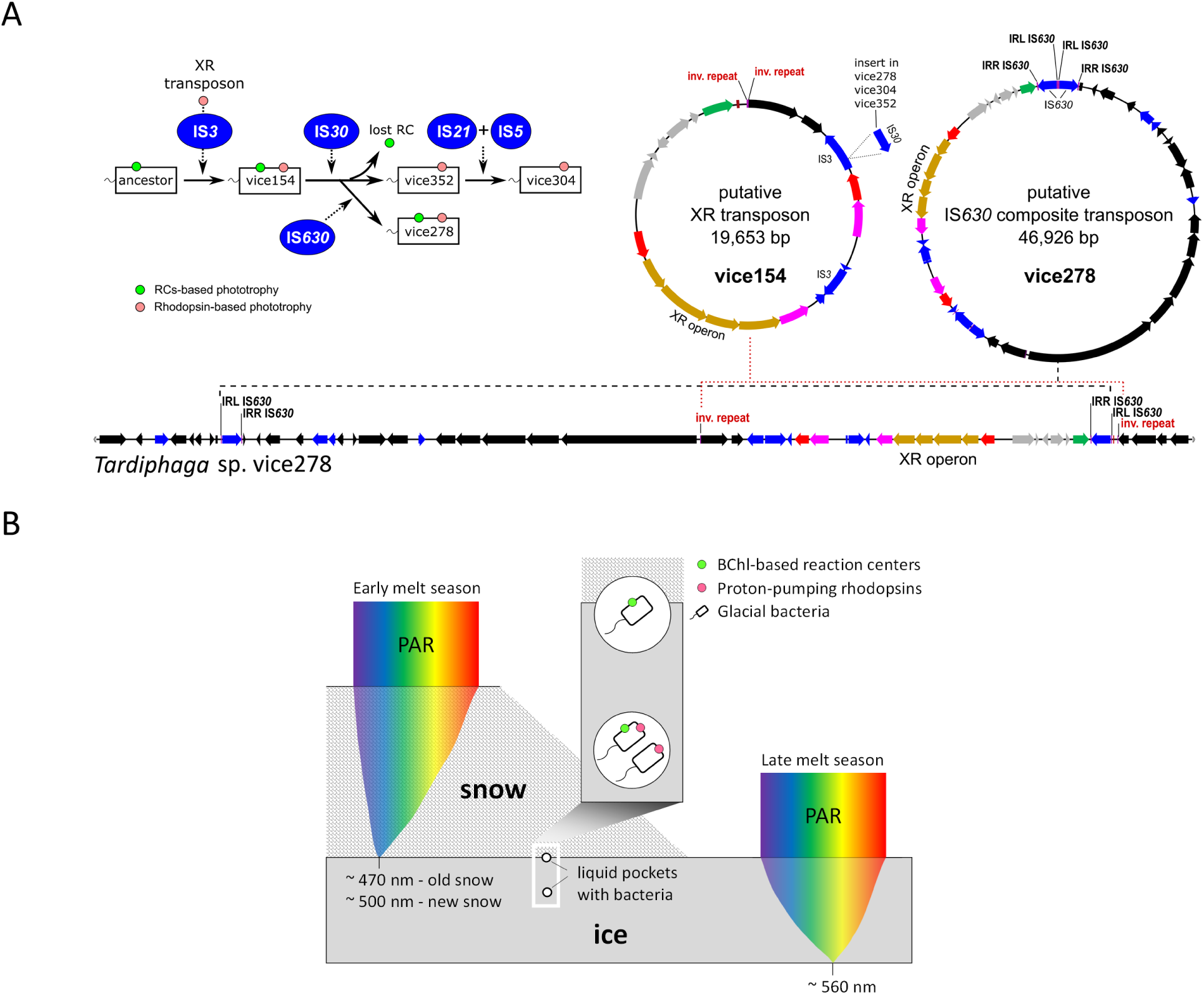
(**A**), hypothetical evolutionary path of *Tardiphaga* isolates from their common ancestor based on IR insertion patterns (left) and two transposons proposed to drive the movement of XR operons (right). The genomic region surrounding XR operon in *Tardiphaga* strain vice278 was shown to highlight the distribution of direct or inverse repeats that are required to form the putative transposons. (**B**), a model for light availability in snow and ice and hypothesized niche partitioning of BChl and rhodopsin‐based dual phototrophy versus only BChl‐ or rhodopsins‐based single phototrophy. The snowpack and icepack are depicted to be of such an ideal thickness that the spectra with the lowest extinction coefficients reach the bottom of snowpack or ice pack. PAR, photosynthetically active radiation. Note that new and old snows have different extinction efficiencies for PAR.

### Ecological importance and wide distribution of potential dual phototrophs

Glacier and ice sheet cover 10% of the land surface of the Earth, hosting an enormous diversity of microbes (19), among which the light‐driven metabolisms by anoxygenic phototrophs have been proposed to have the potential of significantly influencing glacial carbon fluxes (20). The potential dual phototrophy in glacial bacteria described in this study could further amplify the ecological importance of anoxygenic phototrophs in glacial ecosystems.

We assessed the abundance of type‐2 RC‐based and rhodopsin‐based phototrophs in the metagenomes of ES and LF. Type‐2 RC‐based phototrophs in both samples are comparable with LF showing a slightly higher percentage (23.3%) (Fig. 1B). However, rhodopsins‐based phototrophs were almost seven times more abundant in LF than in ES (83.3% vs. 12.2%, Fig. 1B), suggesting that rhodopsins‐based phototrophs may play a more important role in supraglacial environments than type‐2 RC‐based phototrophs, in line with the fact that rhodopsin‐based phototrophy requires less energy and fewer components for assembly and thus probably responds faster to environmental changes than BChl‐based phototrophy.

Dual phototrophy can be particularly advantageous in extreme environments like high Arctic glacial surfaces where phototrophic bacteria may have to exploit all available solar radiation for energy production. Given the advantage of dual phototrophy over single phototrophy in light harvesting, it is unclear as to why PGC was selectively lost in *Tardiphaga* sp. vice304 and vice352. We proposed two theories to explain this phenomenon, i.e. niche differentiation and gene mutation.

XR has a maximum absorption in the green light (9), while BChl and accessory carotenoids in anoxygenic phototrophs mostly absorb blue and infrared light (21). Different light wavelength attenuates differently through snow and ice. Under an ideal condition without impurities, gaps and spatial heterogeneities occurring in snow and ice, blue light tends to reach deepest into the snow (22,23), while green light at approx. 560 nm has the lowest attenuation coefficient within ice (24). Thus, different niches exist in glacial surface in terms of light intensity and quality, which may select for phototrophic bacteria with different preference for light spectra (Fig.2B). The loss of PGC and BChl‐based phototrophy may also occur through random mutation in key PGC genes, caused by, for instance, the activities of transpose gene as seen inside the PGC of vice278 (Fig.1C).

There are nine MAGs reconstructed in this study that contain both *pufM* and rhodopsin genes (Table 1 and Table S2), including five Alphaproteobacteria, three Gammaproteobacteria, and one Gemmatimonadetes, among which all but one were recovered from the LF glacier sample. To test if dual phototrophy is exclusively occurring in supraglacial environment, we further searched public databases (NCBI and ENA, n=215,874, see Supplementary Methods) for bacterial genomes of similar dual phototrophy potential (type‐2 RC and proton‐pumping rhodopsin). We found 3,442 XR/proteorhodopsin‐like and 1,521 *pufM*‐like hits (Table S3). Fifty‐five genomes were predicted to contain both *pufM* and rhodopsin genes (Table S3) with the majority (n=40) belonging to Alphaproteobacteria. Interestingly, there are also four Chloroflexi, three Bacteroidetes, two Deltaproteobacteria, two Gammaproteobacteria, and one Actinobacteria. Given the quality concern on incomplete genomes deposited into public databases (25), it is unclear if there is any composite among these genomes, especially those from Bacteroidetes and Actinobacteria where BChl‐based phototrophy has not yet been reported. The isolation sources of these genomes cover various environment (Table S3), including freshwater, seawater, ground water, hot spring, microbial mat/biofilm, soil, sediment, plant surface, and cryosphere (alpine/polar; n=24), indicating that dual phototrophy is likely present in a wide range of bacteria and in a large variety of natural environments beyond glaciers.

Dual phototrophy is clearly not a metabolic trait that only evolved in glacial bacteria, albeit the four glacial *Tardiphaga* strains in this study and their XR operons both show cryospheric origin (Fig. 1E). We investigated whether these *Tardiphaga* strains have other metabolic traits that may enable them to adapt to the high Arctic glacier environment. Strikingly, all four genomes contain RuBisCO genes, phosphoribulokinase, and soluble methane monooxygenase gene (Table S4), pointing to the metabolic potentials of photoautotrophy and methanotrophy. We have so far failed to grow these *Tardiphaga* strains in liquid culture and neither observed their expression of BChl or XR under tested laboratory conditions (Supplementary Note). Further growth optimization and physiological data are warranted to verify their dual phototrophy and other metabolic potentials and to understand how the two types of phototrophy coordinate in their metabolic networks.

## Materials and Methods available as supporting information

## Acknowledgements

We thank Jørgen Skafte for his excellent technical assistance at the Villum Research Station, Alexandre M. Anesio for discussion, and Niels Bohse Hendriksen for the help during the early stage of this project. We also thank Nupur and Michal Koblížek for their help on our failed experiment of pigment analysis. This work was supported by a Villum Experiment grant (no. 17601) and a Marie Skłodowska‐Curie AIAS‐COFUND fellowship (EU‐FP7, no. 609033) to YZ. Computing time and cost for the bioinformatic analysis were partly supported by Computerome.dk through a fund provided by the Danish e‐Infrastructure Cooperation.

## Author contributions

Y.Z. conceived the study and wrote the paper. Y.Z., X.C., A.M.M, A.Z., L.C.L., Y.L., and L.H.H. contributed to data collection or analysis. T.K.N. analyzed the mobile elements. A.A conducted the rhodopsin sequences classification. All authors read, commented, and approved the paper.

## Competing interests

X.C. is currently employed at BGI Europe A/S, Denmark. The employer did not play a role in the design and practice of this study and neither influenced the writing of this paper in any form. All other authors declare no competing interests.

## Supporting Materials

**Supplementary Methods and Note** Detailed materials and methods in text with a note on the negative results from cultivation of *Tardiphaga* stains in liquid media and colony pigment analysis.

**Table S1** Summary of complete genomes of the four *Tardiphaga* strains isolated from the “Little Firn” glacier in northeast Greenland.

**Table S2** Summary of metagenome‐assembled genomes (MAGs) from the LF and ES metagenomes.

**Table S3** Survey of *pufM* and rhodopsin genes in the genomes deposited into NCBI’s Microbial Genome and ENA’s WGS databases and the list of public genomes containing both *pufM*‐like and rhodopsin‐like genes.

**Table S4** Survey of key functional genes with biogeochemical or energetic importance in the genomes of *Tardiphaga* isolates. The gene list was compiled from the FunGen pipeline (http://fungene.cme.msu.edu/) and the authors’ own collection.

**Figure S1** Photographs showing the sampling site, fieldwork, and visual inspection of a sampled ice block.

**Figure S2** Composition of total bacterial communities in LF and ES at the phylum level (**A**) highlighting Alphaproteobacteria that contains the *Tardiphaga* genus (**B**).

**Figure S3** Annotated genomic regions surrounding PGCs in the *Tardiphaga* strains vice154 and vice278 in comparison to vice304 and vice352 that do not contain a PGC.

**Figure S4** Protein sequence alignments showing the presence of residues in *Tardiphaga*’s xanthorhodopsins that are key to the function as a proton pump.

**Figure S5** Maximum‐likelihood phylogeny of the whole XR operon of *Tardiphaga* isolates and their top tBLASTn hits in NCBI’s genome database.

## Supplementary Methods and Note

### Sampling, cultivation and DNA sequencing

The sampling site was the “Lille Firn” (LF) glacier (81.566° N, 16.363° W) in the Knuths Fjeld area of northeast Greenland, 5.6 km away from the Villum Research Station (VRS). The LF glacier was independent and formed at the lee side of a small hill, surrounded by ∼160 km^2^ land of permafrost. Surface ice was collected on 2 July 2018 using a sharp spear after removal of the top ∼2 m thick snow cover. The sampled ice was processed within 24 hrs in the VRS laboratory. The ice surface was cleaned with running pre‐sterilized and cooled water before melting at 4 °C in sterile whirl‐pak sampling bags (Nasco) for 24‐48 hours. During our fieldwork, the melt season just began and the majority of the whole area was heavily covered by snow. There was a small patch of exposed soil (designated as ES), ca. 50 meters away to the north of the glacier sampling site, where a few kilograms of surface soil (top layer, a few centimeters thick) were collected into a sterile whirl‐pak bag and kept at 4 °C as a comparison site for the following cultivation and metagenomics analyses.

For bacterial cultivation, 100 μL 3.0 μm‐prefiltered meltwater was plated onto 1/5 strength R2A agar (Difco) at the VRS laboratory. The plates were kept aerobically for 8 weeks under room temperature and then screened for bacteriochlorophyll fluorescence using a colony infrared imaging system as described in Zeng et al. (2014). A MALDI‐TOF mass spectrometer (Microflex LT, Bruker Daltonics) was used to generate protein fingerprints for colonies in order to rapidly determine their relatedness at the species level as described previously (Zervas et al., 2019). Genomic DNA of the selected isolates was extracted from cells harvested from two‐week old 1/5 R2A agar plates using the EasyPure bacterial genomic DNA kit (Transgen Biotech, Beijing, China) and were sequenced using both the BGISEQ sequencer (BGI Europe, Denmark) and a MinION device in house following standard protocols as described previously (Zervas et al., 2019). Gap‐free complete genomes were assembled using Unicycler (ver. 0.4.8) in a hybrid mode with default settings (Wick et al., 2017). Genomes were annotated with the NCBI’s prokaryotic genome annotation pipeline. Genome synteny was visualized with the Easyfig program (ver. 2.2.3).

For amplicon and metagenomics analyses, cells in ∼20 L of 3.0 μm‐prefiltered melt ice were collected onto 0.2 μm filters and the total DNA was extracted using the DNeasy PowerWater DNA extraction kit (Qiagen, Germany). Amplicon sequencing of 16S rRNA gene was conducted at BGI Hong Kong targeting the V3‐V4 region and the data were analyzed using the 16S pipeline embedded in the Geneious Prime (Biomatters, New Zealand). The PE reads were first end‐trimmed with quality score set as Q>20 and then merged. Non‐merged reads were discarded and only the high‐quality merged reads were used for the community structure analysis. Total environmental DNA was sequenced on an Illumina NovaSeq platform (BGI Hong Kong). The generated ∼220 G bases of PE reads (150 bp) were end‐trimmed (>Q20, >50 bases long) and assembled using Megahit (ver. 1.1.x; Li et al., 2015) with minimum contig length of 500 bp. There were 948,300 contigs (⩾1 kb; total length, 2.69 G bases) assembled for the LF glacial sample and 2,834,721 contigs (⩾1 kb; total length, 5.63 G bases) for the ES soil sample.

### Binning, database mining and bioinformatics

Binning of metagenome‐assembled contigs was performed using MetaBAT2 with default settings (Kang et al., 2019). Genomic bins were de‐replicated using dRep (Olm et al., 2017) and quality checked with CheckM by following the lineage‐specific workflow (Parks et al., 2015). Only bins of good quality (>50% completeness, <10% contamination; recommended by Bowers et al., 2017) were included for further analysis. Each bin was taxonomically classified using the GTDB‐Tk tool (https://github.com/Ecogenomics/GTDBTk, Parks et al., 2018). Based on the GTDB‐Tk classification results, genomes that have more than 10% of markers with multiple hits were discarded.

The sequences of *pufM* gene (encoding type‐2 reaction center M protein), rhodopsin gene, and the single‐copy gene *recA* were retrieved from the ES and LF metagenome assemblies. The *recA* gene encodes a DNA recombination and repair protein and is commonly used as a phylogenetic biomarker within a similar size range (234 aa) as phototrophy‐related genes (average length: *pufM*, 293 aa; rhodopsin genes, 201 aa). Multiple reference sequences of each gene with wide phylogenetic coverage were used as tBLASTn queries against the ES and LF metagenome assemblies. The cutoff E‐value and minimum identities and coverage for tBLASTn hits were e‐5, 30% sequence identity, and 30% query coverage. The lowest scored hit was confirmed as target gene by BLASTp search against NCBI’s RefSeq protein database. The cleaned tBLASTn results from different reference sequences were pooled for each gene and used for abundance estimates.

To assess the relative abundance of *pufM* and rhodopsin genes in the metagenomes, original reads were mapped onto the assembled contigs using Bowtie2 (Langmead and Salzberg, 2012) and SAMtools (Li et al., 2009). After duplicate reads were removed using the Picard toolkit (https://gatk.broadinstitute.org), mapped reads per target gene (*pufM*, rhodopsin and *recA* genes with their locations on assemblies marked during tBLASTn analysis) were counted using *featureCounts* of the *Subread* package (Liao, et al., 2013) and was further normalized as the number of reads per million reads per kb length of the gene. The relative abundance of phototrophs in the whole community was estimated as the number of total reads mapped to a phototrophic gene divided by the number of total reads mapped to the single‐copy *recA* gene.

Searching for rhodopsin and *pufM* genes in prokaryotic genomes deposited into public databases was carried out as following. First, all retrievable prokaryotic genomes deposited into NCBI’s Microbial Genome database (n=215,874) and ENA’s WGS database (n=227,814) were bulk downloaded from NCBI and ENA’s ftp servers (as of 11 November 2019). Common human‐associated bacteria (NCBI, n=107,120; ENA, n=129,669) were removed from the collection by searching for keywords in *fasta* headers, including *Brucella, Chlamydia, Clostridioides, Clostridium, Corynebacterium, Enterococcus, Escherichia, Haemophilus, Helicobacter, Klebsiella, Listeria, Mycobacterium, Neisseria, Salmonella, Shigella, Staphylococcus, Streptococcus*, and *Yersinia*, and three highly represented species according to the stats in the GTDB genome collection (https://gtdb.ecogenomic.org/stats), i.e. *Acinetobacter baumannii, Pseudomonas aeruginosa*, and *Mycobacteroides abscessus*. Then, the ENA and NCBI genome datasets were merged (non‐redundant, n=108,754) and served as the local BLAST database built with NCBI’s BLAST+ tools (ncbi‐blast‐2.2.18). The *pufM* and *XR* genes of the *Tardiphaga* isolates in this study (Tar_pufM and Tar_XR) and the PR gene of *Pelagibacter* sp. IMCC9063 (Pel_PR) were used as tBLASTn queries. The initial thresholds of 30% sequence identity, 30% query coverage and an E‐value cutoff of e^‐5^Supplementary Methods and Note were applied to filter the tBLASTn results. The tBLASTn hit with the lowest score was further checked by BLAST at the UniProt website until the lowest one was confirmed as target gene.

### Rhodopsin gene classification

Multiple reference rhodopsin genes with wide phylogenetic coverage were used as tBLASTn queries against the ES and LF metagenom‐assembled genomes. The cutoff E‐value and minimum identities and coverage for tBLASTn matches were e‐5, 30% sequence identity, and 30% query coverage. The cleaned tBLASTn results from different reference sequences were pooled for each gene and used for phylogenetic analysis. To reduce the uncertanites when placing short sequences on phylogenetic trees, short sequenes were excluded from phylogenetic analysis but kept in read mapping and abundance estimates. The length cutoffs were <150 amino acids (aa) for rhodopsin genes.

The sequences of rhodopsin genes together with reference sequences were aligned with MUSCLE (Edgar, 2004) and the phylogenetic tree was inferred by following Bulzu et al. (2019). Briefly, the identified rhodopsin sequences (>150 aa, n=775 sequences) were scanned with HMMER against a locally installed Pfam database (version 32) using the script pfam_scan.pl (obtained from the Pfam’s ftp site). As the rhodopsins proved to be composed of both type‐1 (n=657 sequences) and heliorhodopsin (n=128 sequences), we merged them with a previously published database (n=410 sequences) (Bulzu et al., 2019). The rhodopsin sequences (n=1,185) were screened with PREQUAL (Whelan et al., 2018) in order to mask non‐homologous characters, and aligned with the PASTA software (Mirarab et al., 2014) using default settings. A maximum‐likelihood phylogeny was constructed using IQ‐TREE (Nguyen et al., 2015) with the LG+F+G4 substitution model (chosen as the best‐fitting model by ModelFinder) and 1000 ultrafast bootstrap replicates. The phylogenetic tree was used for classification of rhodopsin genes.

### XR operon phylogeny

The translated protein sequences of the six genes in XR operon (XR‐*crtEIBY*‐*brp*) were concatenated and the generated protein sequence (2,060 amino acid sites) was used as the tBLASTn query against NCBI’s RefSeq genome database to search for closely related relatives. All hits that meet the threshold (total query coverage >80%, total sequence identity >50%, and individual gene’s sequence identity >30% and query coverage >50%) were downloaded from NCBI. The protein sequence of each gene in *Tardiphaga*’s XR operon was individually aligned with reference sequences using MUSCLE (Edgar, 2004). All alignments of six genes were concatenated for phylogeny inferrence using FastTree (ver.2.1.12, LG model and gamma approximation with 100 bootstrap replicates; Price et al., 2010) within the Geneious Prime environment.

### Note on the negative results from cultivation of *Tardiphaga* stains in liquid media and colony pigment analysis

Growth of *Tardiphaga* strains in the liquid media was tested on full‐strength R2B (liquid medium version of R2A) and 1/5 R2B for heterotrophic growth and the Rhodospirillaceae medium (DSMZ medium 27 without using L‐cysteiniumchloride and resazurin) that was designed for purple bacterial photoautotrophic growth. The cultivation conditions were 25°C, 16/8 hr light cycle with a 100 W tungsten lamp. For aerobic growth, cotton‐plugged flasks were used with constant shaking at 200 rpm. For anaerobic growth, 125 mL glass serum bottles were used and the test medium was flushed with nitrogen gas and sealed with a rubber septum under a stream of nitrogen gas prior to autoclave at 121°C for 15 min. Sterile syringes were used to inoculate and remove samples and to inject temperature‐sensitive components of the medium. No growth was observed as measured by changes in OD600 during an 11‐week aerobic incubation and almost half a year of anaerobic growth. All four strains were tested and showed negative results.

For pigment analysis, colonies grown on three weeks old 1/5 R2A plates were scraped and the pigment was extracted with 100% methanol. Twenty microliters of the mix was injected into Nexera LC‐40 HPLC system (Shimadzu, Japan; accessed at Michal Koblížek’s group in the Institute of Microbiology CAS, Czech Republic) equipped with Kinetex 2.6 µm C8 100Å column (150 mm × 4.6 mm, Phenomenex) heated at 40°C. A binary solvent system was used: A, 25% 28 mM ammonium acetate + 75% methanol; B, 100% methanol at a constant flow rate 0.8 mL min‐1. BChl *a* peaks and carotenoids were observed at 770 nm and 490 nm respectively. To detect the pigment of salinixanthin that was involved in the light‐harvesting carotenoid antenna of xanthorhodopsin (ref. 9), *Salinibacter ruber* strain DSM 13855 was purchased from DSMZ and used as positive control. No BChl and salinixanthin signals were observed.

## References

1. Pinhassi J, DeLong EF, Béjà O, González JM, Pedrós-Alió C (2016) Marine bacterial and archaeal ion-pumping rhodopsins: genetic diversity, physiology, and ecology. Microbiology and molecular biology reviews 80(4), 929–954.

2. Sabehi G, Loy A, Jung KH, Partha R, Spudich JL, Isaacson T, Hirschberg J, Wagner M, Béja O (2005) New insights into metabolic properties of marine bacteria encoding proteorhodopsins. PLoS Biology, 3(8), p.e273.

3. Kirchman, D. L., & Hanson, T. E. (2013). Bioenergetics of photoheterotrophic bacteria in the oceans. Environmental microbiology reports, 5(2), 188–199.

4. Gómez-Consarnau L, Raven JA, Levine NM, Cutter LS, Wang D, Seegers B et al (2019) Microbial rhodopsins are major contributors to the solar energy captured in the sea. Science Advances 5(8), eaaw8855.

5. Petersen J, Brinkmann H, Bunk B, Michael V, Päuker O, Pradella S (2012) Think pink: photosynthesis, plasmids and the Roseobacter clade. Environmental microbiology, 14:2661–2672.

6. Brinkmann H, Göker M, Koblížek M, Wagner-Döbler I, Petersen J (2018) Horizontal operon transfer, plasmids, and the evolution of photosynthesis in Rhodobacteraceae. The ISME Journal 12: 1994.

7. Liu Y, Zheng Q, Lin W, Jiao N (2019) Characteristics and Evolutionary analysis of Photosynthetic Gene Clusters on Extrachromosomal Replicons: from streamlined plasmids to chromids. mSystems 4:e00358–19.

8. Yutin N, Koonin EV (2012) Proteorhodopsin genes in giant viruses. Biology direct 7(1), 34.

9. Balashov SP, Imasheva ES, Boichenko, VA, Antón J, Wang JM, Lanyi JK (2005) Xanthorhodopsin: a proton pump with a light-harvesting carotenoid antenna. Science 309: 2061–2064.

10. Thiel V, Hügler M, Ward DM, Bryant DA (2017) The dark side of the mushroom spring microbial mat: life in the shadow of chlorophototrophs. II. Metabolic functions of abundant community members predicted from metagenomic analyses. Frontiers in microbiology, 8, 943.

11. Tank M, Thiel V, Ward DM, Bryant DA (2017) A panoply of phototrophs: an overview of the thermophilic chlorophototrophs of the microbial mats of alkaline siliceous hot springs in Yellowstone National Park, WY, USA. In Modern topics in the phototrophic prokaryotes (pp. 87–137). Springer, Cham.

12. Zeng, Y., Feng, F., Medová, H., Dean, J., & Koblížek, M. (2014). Functional type 2 photosynthetic reaction centers found in the rare bacterial phylum Gemmatimonadetes. Proceedings of the National Academy of Sciences, 111(21), 7795–7800.

13. De Meyer SE, Coorevits A, Willems A (2012) *Tardiphaga robiniae* gen. nov., sp. nov., a new genus in the family Bradyrhizobiaceae isolated from *Robinia pseudoacacia* in Flanders (Belgium). Systematic and applied microbiology 35(4), 205–214.

14. Boyd, E. F., Almagro-Moreno, S., & Parent, M. A. (2009). Genomic islands are dynamic, ancient integrative elements in bacterial evolution. Trends in microbiology, 17(2), 47–53.

15. Béja, O., Aravind, L., Koonin, E. V., Suzuki, M. T., Hadd, A., Nguyen, L. P., … & Spudich, E. N. (2000). Bacterial rhodopsin: evidence for a new type of phototrophy in the sea. Science, 289(5486), 1902–1906.

16. Frigaard NU, Martinez A, Mincer TJ, DeLong EF (2006) Proteorhodopsin lateral gene transfer between marine planktonic Bacteria and Archaea. Nature 439:847.

17. Ugalde, J. A., Podell, S., Narasingarao, P., & Allen, E. E. (2011). Xenorhodopsins, an enigmatic new class of microbial rhodopsins horizontally transferred between archaea and bacteria. Biology direct, 6(1), 52.

18. Ward, L. M., Hemp, J., Shih, P. M., McGlynn, S. E., & Fischer, W. W. (2018). Evolution of phototrophy in the Chloroflexi phylum driven by horizontal gene transfer. Frontiers in microbiology, 9, 260.

19. Anesio, A. M., & Laybourn-Parry, J. (2012). Glaciers and ice sheets as a biome. Trends in ecology & evolution, 27(4), 219–225.

20. Ferrera, I., Sánchez, O., Kolářová, E., Koblížek, M., & Gasol, J. M. (2017). Light enhances the growth rates of natural populations of aerobic anoxygenic phototrophic bacteria. The ISME journal, 11(10), 2391–2393.

21. Stomp, M., Huisman, J., Stal, L. J., & Matthijs, H. C. (2007). Colorful niches of phototrophic microorganisms shaped by vibrations of the water molecule. The ISME journal, 1(4), 271–282.

22. Grenfell, T. C., & Maykut, G. A. (1977). The optical properties of ice and snow in the Arctic Basin. Journal of Glaciology, 18(80), 445–463.

23. Hancke, K., Lund - Hansen, L. C., Lamare, M. L., Højlund Pedersen, S., King, M. D., Andersen, P., & Sorrell, B. K. (2018). Extreme low light requirement for algae growth underneath sea ice: A case study from Station Nord, NE Greenland. Journal of Geophysical Research: Oceans, 123(2), 985–1000.

24. Perovich, D. K., Roesler, C. S., & Pegau, W. S. (1998). Variability in Arctic sea ice optical properties. Journal of Geophysical Research: Oceans, 103(C1), 1193–1208.

25. Shaiber, A., & Eren, A. M. (2019). Composite metagenome-assembled genomes reduce the quality of public genome repositories. mBio, 10(3), e00725–19.

## References

Bowers RM, Kyrpides NC, Stepanauskas R, Harmon-Smith M, Doud D, Reddy TB, Schulz F, Jarett J, Rivers AR, Eloe-Fadrosh EA, Tringe SG (2017) Minimum information about a single amplified genome (MISAG) and a metagenome-assembled genome (MIMAG) of bacteria and archaea. Nature biotechnology 35(8):725.

Bulzu, P. A., Andrei, A. Ş., Salcher, M. M., Mehrshad, M., Inoue, K., Kandori, H., … & Banciu, H. L. (2019). Casting light on Asgardarchaeota metabolism in a sunlit microoxic niche. Nature microbiology, 4(7), 1129–1137.

Edgar, R. C. (2004). MUSCLE: a multiple sequence alignment method with reduced time and space complexity. BMC bioinformatics, 5(1), 113.

Kang D, Li F, Kirton ES, Thomas A, Egan RS, An H, Wang Z. (2019) MetaBAT 2: an adaptive binning algorithm for robust and efficient genome reconstruction from metagenome assemblies. PeerJ 7:e27522v1.

Langmead, B., & Salzberg, S. L. (2012). Fast gapped-read alignment with Bowtie 2. Nature methods, 9(4), 357.

Li, H., Handsaker, B., Wysoker, A., Fennell, T., Ruan, J., Homer, N., … & Durbin, R. (2009). The sequence alignment/map format and SAMtools. Bioinformatics, 25(16), 2078–2079.

Liao, Y., Smyth, G. K., & Shi, W. (2013). The Subread aligner: fast, accurate and scalable read mapping by seed-and-vote. Nucleic acids research, 41(10), e108–e108.

Mirarab, S., Nguyen, N., Guo, S., Wang, L. S., Kim, J., & Warnow, T. (2015). PASTA: ultra-large multiple sequence alignment for nucleotide and amino-acid sequences. Journal of Computational Biology, 22(5), 377–386.

Nguyen, L. T., Schmidt, H. A., Von Haeseler, A., & Minh, B. Q. (2015). IQ-TREE: a fast and effective stochastic algorithm for estimating maximum-likelihood phylogenies. Molecular biology and evolution, 32(1), 268–274.

Olm MR, Brown CT, Brooks B, Banfield JF (2017) dRep: a tool for fast and accurate genomic comparisons that enables improved genome recovery from metagenomes through de-replication. The ISME journal, 11(12):2864.

Parks DH, Chuvochina M, Waite DW, Rinke C, Skarshewski A, Chaumeil PA, Hugenholtz P (2018) A standardized bacterial taxonomy based on genome phylogeny substantially revises the tree of life. Nature Biotechnology, 36:996–1004.

Parks DH, Imelfort M, Skennerton CT, Hugenholtz P, Tyson GW (2015) CheckM: assessing the quality of microbial genomes recovered from isolates, single cells, and metagenomes. Genome research, 25(7):1043–55.

Price, M. N., Dehal, P. S., & Arkin, A. P. (2010). FastTree 2–approximately maximum-likelihood trees for large alignments. PloS one, 5(3).

Wick RR, Judd LM, Gorrie CL, Holt KE (2017) Unicycler: resolving bacterial genome assemblies from short and long sequencing reads. PLoS Computational Biology, 13(6), e1005595.

Zervas, A., Zeng, Y., Madsen, A. M., & Hansen, L. H. (2019). Genomics of Aerobic Photoheterotrophs in Wheat Phyllosphere Reveals Divergent Evolutionary Patterns of Photosynthetic Genes in *Methylobacterium* spp. Genome biology and evolution, 11(10), 2895–2908.

