## Supplementary figures and images for "Potential Rhodopsin and Bacteriochlorophyll-Based Dual Phototrophy in a High Arctic Glacier"

### Figure S1

#### Figure S1

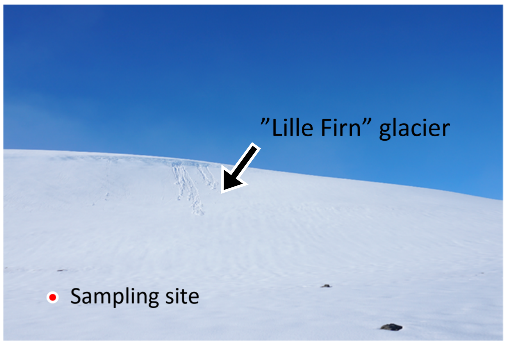


Sampled on 2 July 2018

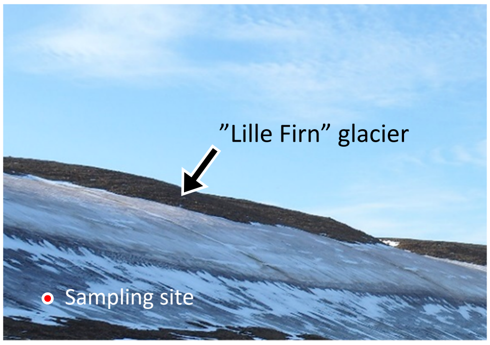


Photographed in August 2017

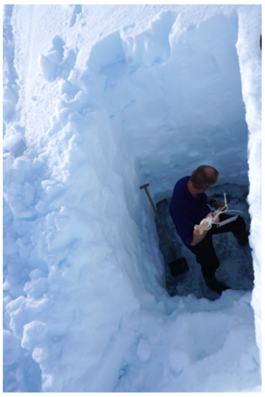

Fieldwork on 2 July, 2018

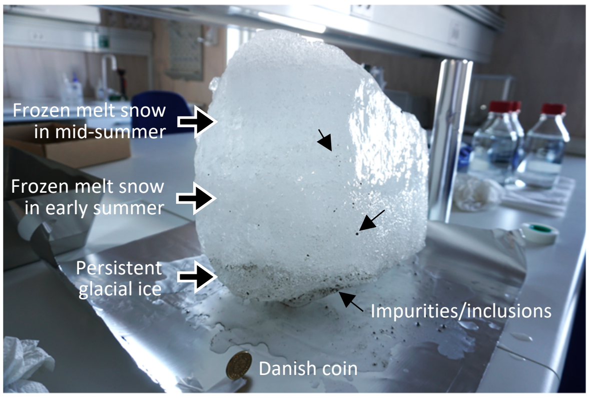


Sampled ice

### Figure S2

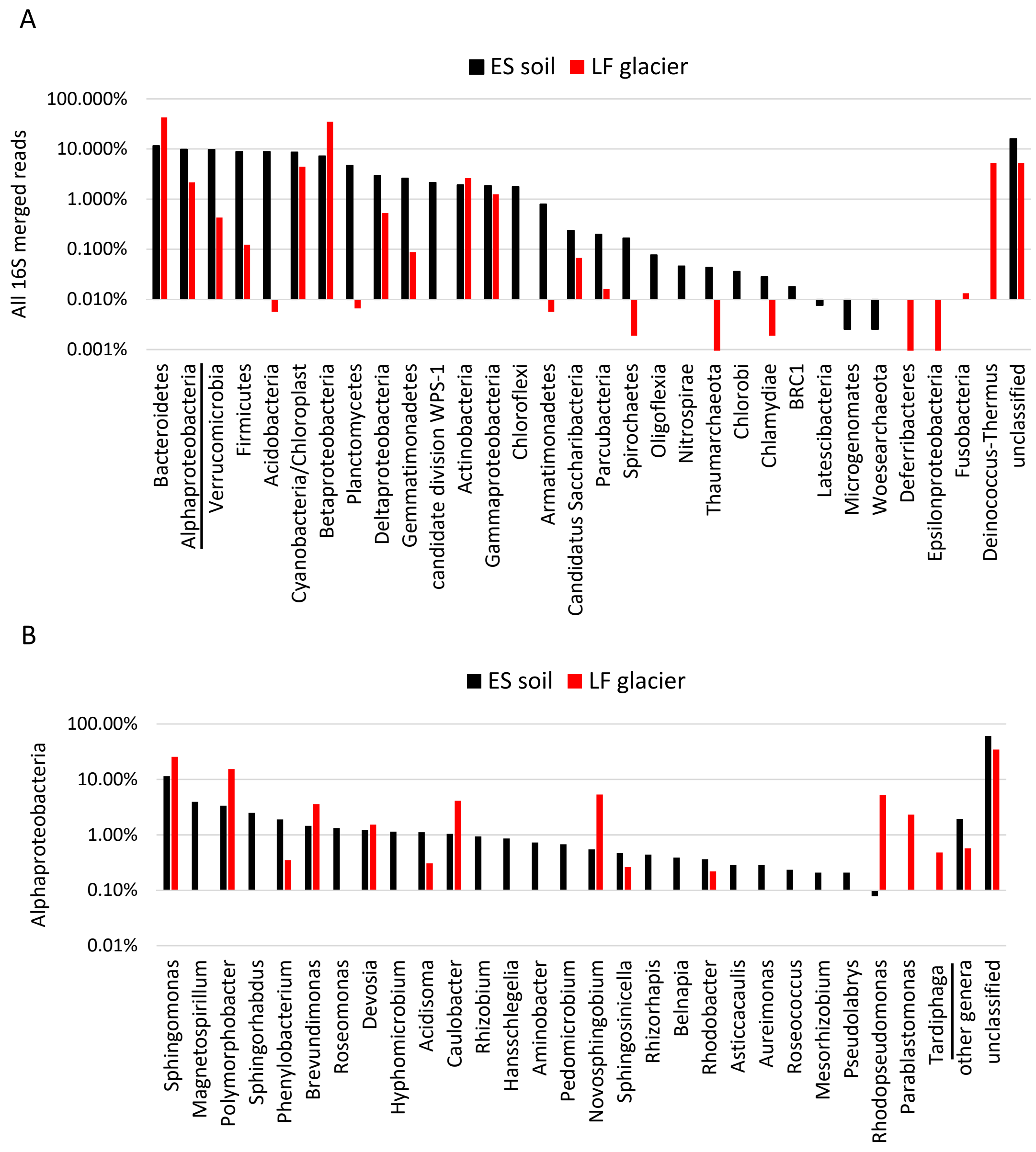

### Figure S3

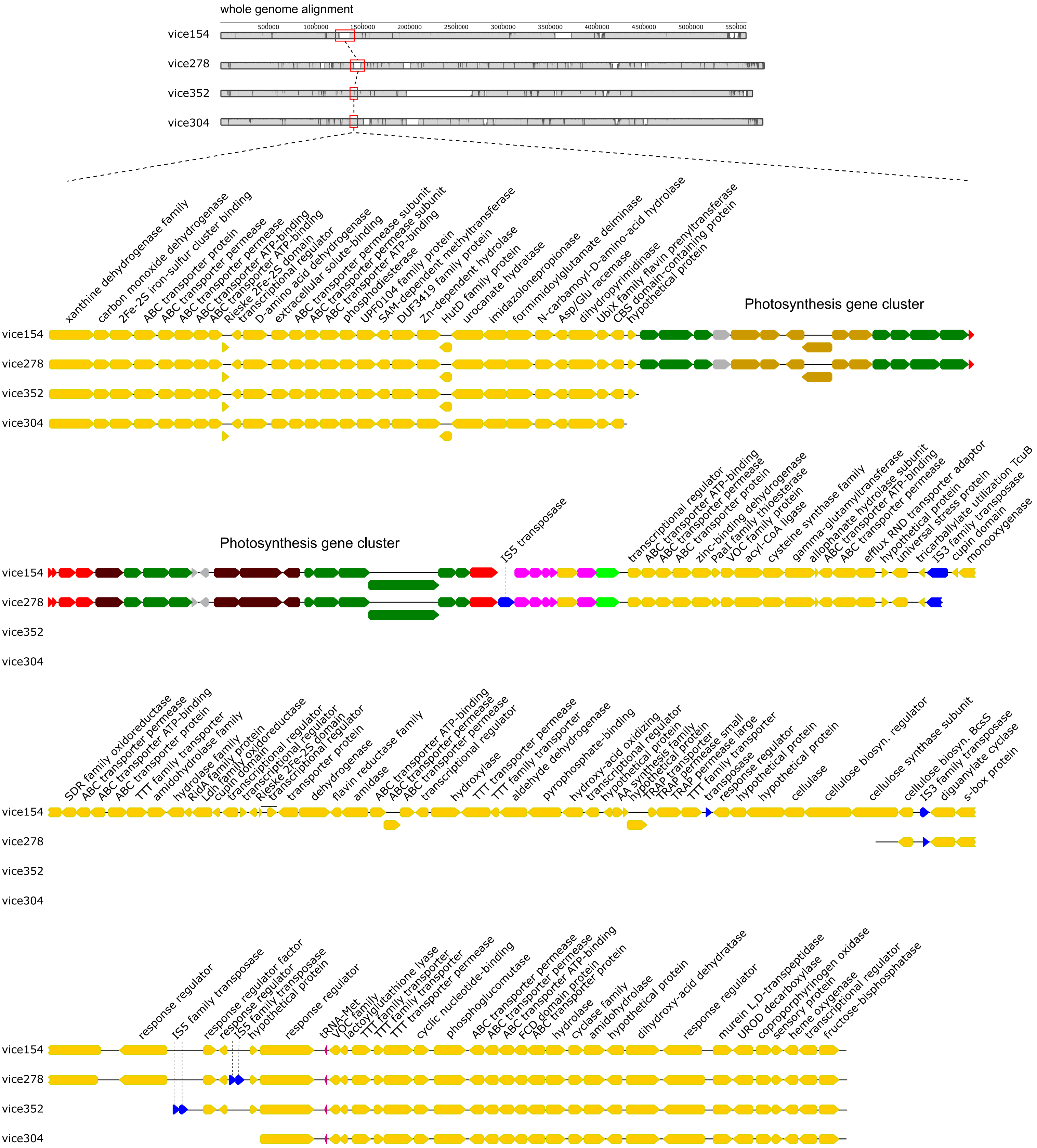

### Figure S4

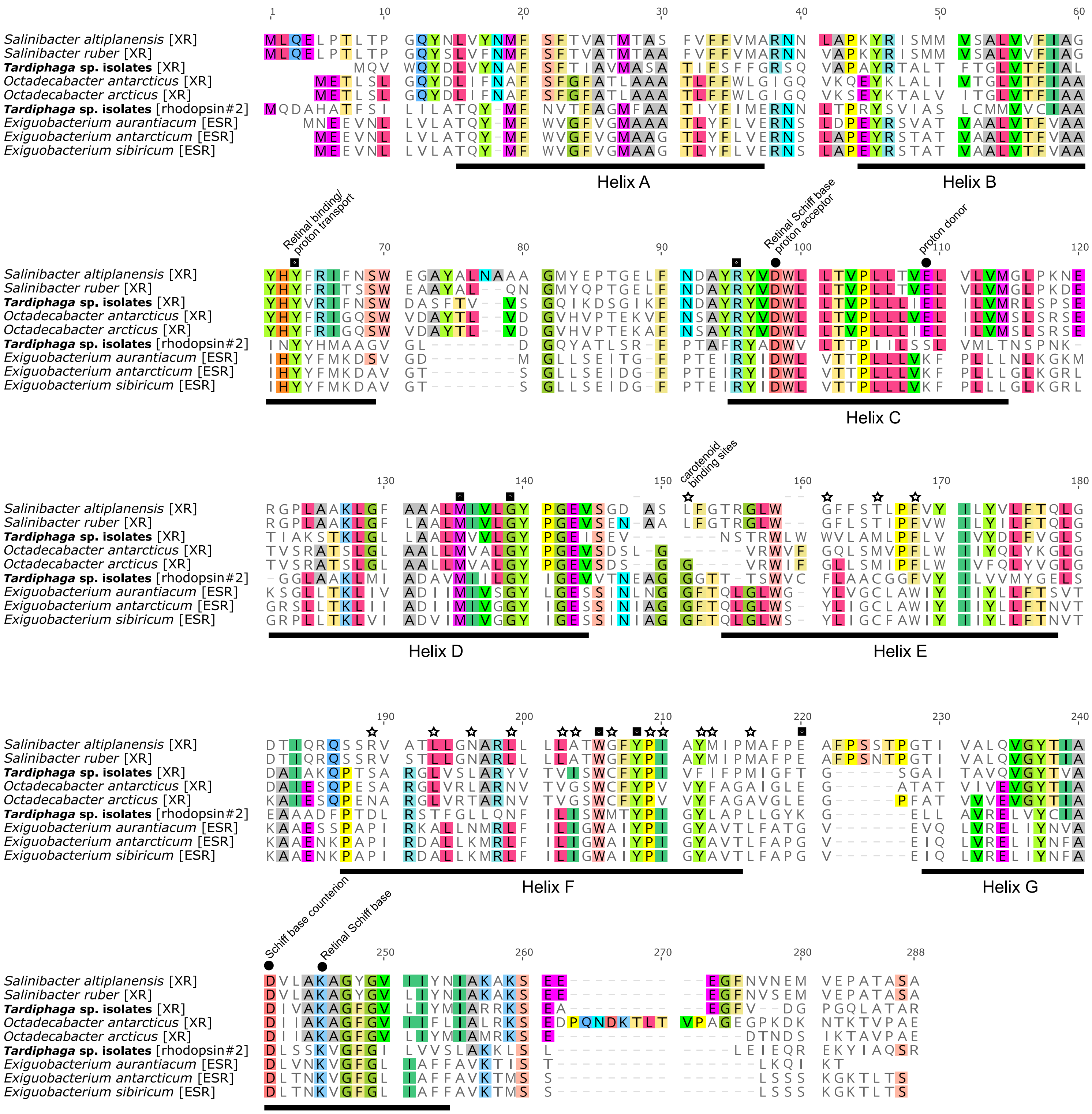

### Figure S5

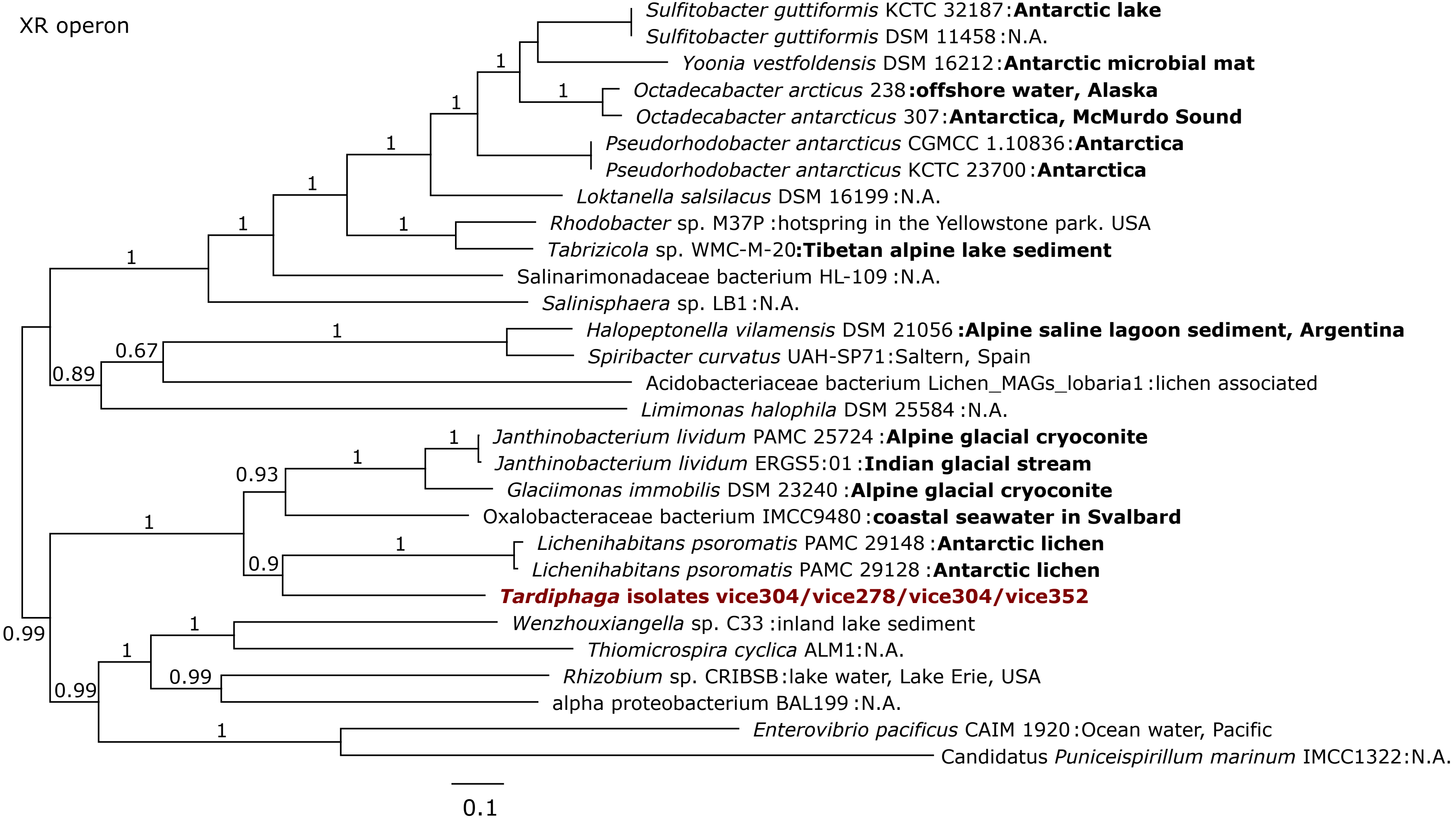
