## Supplementary material for "Potential Rhodopsin and Bacteriochlorophyll-Based Dual Phototrophy in a High Arctic Glacier": Table S1

**Table S1** Summary of the complete genomes of the four *Tardiphaga* strains isolated from the "Little Firn" glacier in northeast Greenland. PGC, photosynthesis gene cluster; XR, xanthorhodopsin. CDS, coding sequence.

| Isolate | Genome Size (bp) | GC content | Number of genes | | | | | | PGC | XR | GenBank accession no. |
|---|---|---|---|---|---|---|---|---|---|---|---|
| | | | rRNA operon | tRNA | Protein CDSs | Trans-posase | Phage-related | Recombinase /integrase | | | |
| vice154 | 5,609,510 | 63.59% | 2 | 51 | 5,001 | 109 | 18 | 16 | + | + | CP041399 |
| vice278 | 5,806,756 | 63.36% | 2 | 53 | 5,196 | 222 | 38 | 28 | + | + | CP041400 |
| vice304 | 5,788,809 | 63.28% | 2 | 53 | 5,192 | 207 | 28 | 33 | - | + | CP041402 |
| vice352 | 5,678,525 | 63.37% | 2 | 52 | 5,094 | 180 | 41 | 26 | - | + | CP041401 |
