## Supplementary material for "Potential Rhodopsin and Bacteriochlorophyll-Based Dual Phototrophy in a High Arctic Glacier": Table S4

Table S4 Survey of key functional genes with biogeochemical or energetic importance in the genomes of *Tardiphaga* isolates. The gene list was complied from the FunGen pipeline (http://fungene.cme.msu.edu/) and the authors' own collection.

| **Category** | **Gene** | **Enzyme** | **vice154** | **vice278** | **vice304** | **vice352** |
|---|---|---|---|---|---|---|
| C metabolism | scd2 | esterase / lipase | ● | ● | ● | ● |
| C metabolism | xylA | xylose isomerase | ○ | ○ | ○ | ○ |
| One carbon metabolism | cooS | carbon monoxide dehydrogenase | ○ | ○ | ○ | ○ |
| One carbon metabolism | pmoA | particulate methane monooxygenase A-subunit | ○ | ○ | ○ | ○ |
| One carbon metabolism | pxmA1 | particulate methane monooxygenase beta subunit | ○ | ○ | ○ | ○ |
| One carbon metabolism | prk | phosphoribulokinase | ● | ● | ● | ● |
| One carbon metabolism | cbbL | ribulose-bisphosphate carboxylase large subunit | ● | ● | ● | ● |
| One carbon metabolism | cbbM | ribulose-bisphosphate carboxylase small subunit | ● | ● | ● | ● |
| One carbon metabolism | smmo | soluble methane monooxygenase | ● | ● | ● | ● |
| N metabolism | amiE | aliphatic amidase | ● | ● | ● | ● |
| N metabolism | amoA | ammonia monooxygenase subunit A | ○ | ○ | ○ | ○ |
| N metabolism | ansA | asparaginase | ○ | ○ | ○ | ○ |
| N metabolism | aspA | aspartate ammonia-lyase | ○ | ○ | ○ | ○ |
| N metabolism | glsA | glutaminase | ● | ● | ● | ● |
| N metabolism | hutH | histidine ammonia-lyase | ○ | ○ | ○ | ○ |
| N metabolism | norB | nitric oxide reductase | ○ | ○ | ○ | ○ |
| N metabolism | p450nor | nitric oxide reductase (NAD(P), nitrous oxide-forming) | ○ | ○ | ○ | ○ |
| N metabolism | nir | nitrite reductase | ● | ● | ● | ● |
| N metabolism | nifH | nitrogenase iron protein | ○ | ○ | ○ | ○ |
| N metabolism | anfD | nitrogenase iron-iron protein, alpha chain | ○ | ○ | ○ | ○ |
| N metabolism | nifD | nitrogenase molybdenum-iron protein subunit alpha | ○ | ○ | ○ | ○ |
| N metabolism | vnfD | nitrogenase vanadium-iron protein alpha chain | ○ | ○ | ○ | ○ |
| N metabolism | nosZ | nitrous oxide reductase catalytic subunit | ○ | ○ | ○ | ○ |
| N metabolism | napA | periplasmic nitrate reductase catalytic subunit | ● | ● | ● | ● |
| N metabolism | napE | periplasmic nitrate reductase subunit | ● | ● | ● | ● |
| N metabolism | apr | serine 3-dehydrogenase | ○ | ○ | ○ | ○ |
| N metabolism | ureC | urease | ● | ● | ● | ● |
| P metabolism | phy | 4-phytase | ● | ● | ● | ● |
| P metabolism | alp | alkaline phosphatase | ● | ● | ● | ● |
| P metabolism | phnX | phosphonoacetaldehyde dehydrogenase | ○ | ○ | ○ | ○ |
| P metabolism | PPK | polyphosphate kinase | ● | ● | ● | ● |
| S metabolism | dsrA | dissimilatory sulfite reductase alpha subunit | ● | ● | ● | ● |
| S metabolism | soxB | sulfur oxidation protein / thiosulfohydrolase | ● | ● | ● | ● |
| Plant material utilization | cbh1 | cellobiohydrolase | ○ | ○ | ○ | ○ |
| Plant material utilization | chiA | chitinase | ○ | ○ | ○ | ○ |
| Plant material utilization | chb | chitobiase | ○ | ○ | ○ | ○ |
| Plant material utilization | ligE | glutathione S-transferase | ○ | ○ | ○ | ○ |
| Plant material utilization | lcc | laccase | ○ | ○ | ○ | ○ |
| Plant material utilization | lip | lignin peroxidase | ○ | ○ | ○ | ○ |
| Plant material utilization | appA | phosphoanhydride phosphohydrolase | ○ | ○ | ○ | ○ |
| Protein catabolism | npr | bacillolysin | ○ | ○ | ○ | ○ |
| Protein catabolism | sub | subtilisin | ○ | ○ | ○ | ○ |
| Protein catabolism | trp | trypsin | ● | ● | ● | ● |
| Phototrophy | acsF | magnesium-protoporphyrin IX monomethyl ester aerobic oxidative cyclase | ● | ● | ● | ● |
| Phototrophy | pufL | photosynthetic reaction center subunit L | ● | ● | ● | ● |
| Phototrophy | pufM | photosynthetic reaction center subunit M | ● | ● | ● | ● |
| Phototrophy | rho | rhodopsin | ● | ● | ● | ● |
| Oxygen fluctuation | ctaD | cytochrome c oxidase subunit, aa3-type, low affinity | ● | ● | ● | ● |
| Oxygen fluctuation | cydA | cytochrome bd-I ubiquinol oxidase subunit 1, bd-type, high affinity | ● | ● | ● | ● |
| Oxygen fluctuation | cyoB | cytochrome bo(3) ubiquinol oxidase subunit, bo3-type, low affinity | ● | ● | ● | ● |
| Oxygen fluctuation | cbaA | cytochrome c oxidase subunit, ba3-type, high affinity | ● | ● | ● | ● |
| Oxygen fluctuation | fixN | cytochrome c oxidase subunit, cbb3-type, high affinity | ● | ● | ● | ● |
| Oxygen fluctuation | ccoN | cytochrome c oxidase subunit, cbb3-type, high affinity | ○ | ○ | ○ | ○ |
| Sense and regulation | PAP | serine/threonine protein phosphatase | ● | ○ | ● | ● |
| Sense and regulation | PTP | tyrosine protein phosphatase | ● | ● | ● | ● |
| Other - acetate metabolism | acs | acetyl-CoA synthetase | ○ | ○ | ○ | ○ |
| Other - animal material utilization | col | collagenase | ○ | ○ | ○ | ○ |
| Other - arginine catabolism | rocF | arginase | ● | ● | ● | ● |
| Other - folate metabolism | cpg | glutamate carboxypeptidase | ○ | ○ | ○ | ○ |
| Other - glycan metabolism | nag3 | beta-hexosaminidase | ○ | ○ | ○ | ○ |
| Other - H metabolism | hydA | iron hydrogenase | ○ | ○ | ○ | ○ |
| Other - $H_2O_2$ generation | glx | glyoxaloxidase 1 | ○ | ○ | ○ | ○ |
| Other - methanogenesis | mcrA | methyl-coenzyme M reductase, alpha subunit | ○ | ○ | ○ | ○ |
| Other - oligosaccharides metabolism | exc1 | beta-N-acetylglucosaminidase | ○ | ○ | ○ | ○ |
| Other - phenol degradation | ppo | phenoloxidase | ○ | ○ | ○ | ○ |
| Other - purine metabolism | add | adenosine deaminase | ○ | ○ | ○ | ○ |
